# Need for speed: Examining protein behaviour during cryoEM grid preparation at different timescales

**DOI:** 10.1101/2020.05.14.095372

**Authors:** David P. Klebl, Molly S. C. Gravett, Dimitrios Kontziampasis, David J. Wright, Robin S. Bon, Diana Monteiro, Martin Trebbin, Frank Sobott, Howard D. White, Michele Darrow, Rebecca F. Thompson, Stephen P. Muench

## Abstract

A host of new technologies are under development to improve the quality and reproducibility of cryoEM grid preparation. Here we have systematically investigated the preparation of three macromolecular complexes using three different vitrification devices (Vitrobot™, chameleon and a time-resolved cryoEM device) on various timescales, including grids made within 6 ms, (the fastest reported to date), to interrogate particle behaviour at the air-water interface for different timepoints. Results demonstrate that different macromolecular complexes can respond to the thin film environment formed during cryoEM sample preparation in highly variable ways, shedding light on why cryoEM sample preparation can be difficult to optimise. We demonstrate that reducing time between sample application and vitrification is just one tool to improve cryoEM grid quality, but that it is unlikely to be a generic ‘silver bullet’ for improving the quality of every cryoEM sample preparation.

## Introduction

Single particle cryo-electron microscopy (cryoEM) has emerged as a major structural biology technique during the last decade (Kuehlbrandt 2014). While refined data processing software (Fernandez-Leiro & Scheres 2017; Punjani et al. 2017; la Rosa-Trevin et al. 2016) and automated data acquisition (Thompson et al. 2019) have streamlined the technique, sample preparation remains a major bottleneck for many projects. Single particle cryo-EM sample preparation requires the specimen to be spread as a thin (≤ 20 to 80 nm) liquid film (Rice et al. 2018) before being rapidly vitrified by plunging into a cryogen liquid such as ethane (Dubochet & Lepault 1984). The formation of this thin film has commonly been achieved by applying a relatively large sample volume (3-4 μL) to a cryoEM grid, and then blotting away excess liquid with filter paper. The cryoEM grid, a 3 mm diameter metal (commonly copper) disk with square windows, has a support layer (typically amorphous carbon), with small, usually circular perforations (~1-2 μm diameter) in a regular array. The typical blotting process removes almost all of the liquid applied to the grid, leaving a thin film of sample suspended across the holes in the support where imaging can occur. This procedure was pioneered over 30 years ago by Dubochet and colleagues (Dubochet & Lepault 1984).

Formation of a thin film using blotting paper followed by vitrification can be achieved through manual and home-built devices, as well as using commercially available devices such as the Vitrobot™ (Thermo Fisher Scientific), EM GP (Leica Microsystems) and CP3 (Gatan), where the general concept remains the same as when the method was first conceived. While there can be problems with reproducibility of thin film formation through a blotting approach it is undeniably successful, resulting in its application to a broad range of specimens, and it has consequently come to underpin the vast majority of single particle structures to date.

Over the years, and across different fields of research, it has been shown that the air-water interface (AWI) can be a hostile environment for proteins and macromolecular complexes (Glaeser & Han 2017; Zhao & Cieplak 2017; Gerhardt et al. 2014; Wiesbauer et al. 2013). In a typical cryoEM grid preparation both sides of the thin film are exposed to the AWI, creating a very high surface area to volume ratio. Blotting and plunging into cryogens usually takes seconds, during which time the sample, can come into contact the AWI 100-1000s of times. Macromolecular complexes and proteins can interact preferentially with and/or denature (either fully or partially) on exposure to the AWI (Taylor, K.A. & Glaeser, R.M., 2008, D’Imprima et al. 2019).

A recent systematic study of particle localisation on cryo-EM grids prepared with traditional blotting methods by Noble *et al.* has shown that ~90 % of the 46 samples analysed associate with the AWI (Noble, Dandey, et al. 2018), demonstrating the vast majority of specimens have the potential to be perturbed by the AWI. Recent advancements have led to a greater awareness of variables that can be changed to alter the distribution and behaviour of particles on a cryoEM grid. These include the use of grid supports made of different materials such as carbon or gold (Russo & Passmore 2016), the use of continuous support films (Hurdiss et al. 2016; Han:2020hq Russo & Passmore 2014), affinity grids (Han et al. 2012), the addition of detergents or surfactants(Chen et al. 2019), or reducing the time between sample application and vitrification (Noble, Wei, et al. 2018). All of these approaches are linked by a common theme – they either sequester particles away from an AWI or they modulate the properties of the AWI by adjusting chemical properties and surface tension of the liquid film (Glaeser & Han 2017).

The grid making process is currently a major focus in the cryoEM field with a number of approaches in various stages of development, all seeking to improve access, quality, and/or reproducibility of cryoEM sample preparation. The Spotiton system uses an inkjet piezo dispenser to directly deposit samples onto self-wicking grids to create a thin film, and is currently undergoing commercialisation (chameleon, SPT Labtech, formerly TTP Labtech) (Razinkov et al. 2016; Wei et al. 2018; Dandey et al. 2018). An alternative open-source approach, the “Shake-it-off” uses an off-the-shelf ultrasonic humidifier to spray small sample volumes onto an EM grid and offers a low-cost solution to grid preparation (Rubinstein et al. 2019). The cryoWriter system uses a microcapillary to deposit sample directly on the grid, enabling direct purification and vitrification from low volumes of lysate (Arnold et al. 2017; Schmidli et al. 2019). The Vitrojet (CryoSol) uses a pin printing system to deposit small volumes of sample on the surface of a grid to directly create a thin film in a controlled manner, followed by vitrification with jets of cryogen (Ravelli et al. 2019). Finally, microfluidic spraying devices such as the **t**ime-resolved cryo-**E**M **d**evice (TED) enable fast dispense-to-plunge times (Kontziampasis et al. 2019) but require larger sample volumes. In this study we focus on the behaviour of particles prepared for cryoEM using the Vitrobot™ Mk IV, TED, and chameleon. Since each of these sample preparation devices exposes particles to different environments, forces, and timescales, we will briefly describe the specifics of each device.

The Vitrobot™ involves the application of 3-4 μL of sample volume onto an EM grid held in a temperature- and humidity-controlled chamber. Subsequently it is blotted between two sheets of filter paper, for 3-10 seconds, removing the vast majority of the sample volume, before the blotting paper is withdrawn and the sample is plunged into the cryogen. Grids can be prepared on a timescale of 5-15 seconds from sample application, and typical grids will have a gradient containing some areas that are too thick and some that are too thin, with a large number of suitable grid squares for imaging (Figure 1A) (Thompson et al. 2019). While this device has been used to successfully vitrify a wide range of specimens, there is evidence that the irregular pattern of fibres in the filter paper causes non-uniform alterations in surface-to-volume ratio across the grid, and this may be a root cause of the irreproducibility often reported for blotting paper-based vitrification techniques, as well as being detrimental to samples (Armstrong et al. 2020).

**Figure 1.**
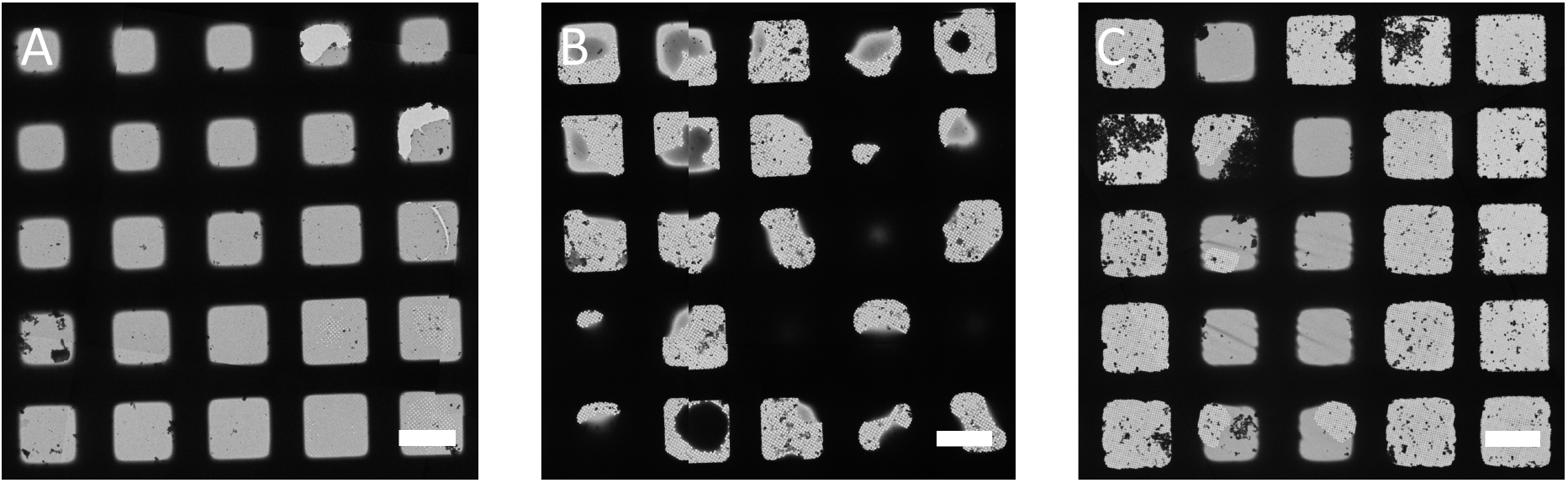
Example low magnification images of grids prepared using different vitrification devices. Comparison of typical results for (A) Vitrobot™, (B) TED and (C) Chameleon (scale bar 50 μm) as imaged by cryoEM.

The TED was primarily designed to perform time-resolved experiments by rapidly mixing constituents before vitrification on the millisecond timescale. However, in this study we only make use of its ability to deposit a single sample and vitrify on a very fast timescale (≥ 6 ms) (Kontziampasis et al. 2019). A conventional EM grid is placed on a plunging arm which has an adjustable speed within a high humidity chamber at room temperature. The liquid system (syringes, tubing and nozzle) is then equilibrated with ~40 μL sample, which is deposited by spraying directly onto the grid as it plunges into the cryogen. A typical experiment requires between 4 and 32 μL of sample volume per grid, depending mainly on the liquid flow rate. Exposure time to the AWI is determined by the time of flight for the spray droplets (from nozzle to grid) and the grid plunge time (from spray to ethane). A typical grid has a random droplet pattern, with some thick regions corresponding to the centre of a droplet, and thinner edges (which sometimes cover ~½ of a grid square) where the ice is sufficiently thin for imaging (Figure 1B). With the current design, dispense-to-plunge times can be set from 6 ms to seconds.

The chameleon is a fully automated instrument which dispenses controlled droplets onto a self-wicking grid as it plunges into the cryogen. Self-wicking grids and 5 μL of sample are manually placed into the instrument as input. Workflows guide the user through system set-up, preparation of grids and system clean-up and reporting. Automated assessment of wicking and visual inspection together provide a quality control step prior to cryoEM allowing the routine preparation of grids with optimal ice thickness. Dispense-to-plunge times range from 54 ms to a few seconds with typical times in the range of 100-250 ms. A typical grid contains a stripe of approximately 20-40 grid squares with desired ice thickness (Figure 1C).

For this study, we have examined the behaviour of three protein systems, apoferritin (480 kDa, O symmetry), mitochondrial chaperone heat shock protein family D member 1 (HSPD1: mtHSPD1) (408 kDa, C7 symmetry) and *Escherichia coli (E. coli)* ribosome (30S, 50S, 70S, all C1 symmetry). Apoferritin was chosen as it is a common test specimen in cryoEM, HSPD1 because when prepared using standard cryoEM methods it adopts an extremely preferred orientation, and ribosomes because they are considered to be a very robust macromolecular complex, and are also asymmetric, unlike the other two specimens.

## Results

### Partitioning of particles to the AWI

The speed of grid making has been reported to influence the particle distribution at the AWI, with ~100 ms showing a change in partitioning and angular orientation relative to slower speeds (Noble, Dandey, et al. 2018). We used cryo-electron tomography (cryoET) to investigate differences in particle partitioning in the thin ice layer at different time points for various macromolecular complexes, using the Vitrobot™, TED, and chameleon (Figure 2A). Areas for tomogram acquisition were selected without prior investigation of particle distribution in that area, and based upon ice thicknesses that would be deemed most suitable for data collection. We classified particles as partitioned to the AWI based on either a 10 nm or 20 nm distance from the AWI. For all three specimen types, on blotted Vitrobot™ grids (Figure 2B) the majority of particles resided at the AWI, consistent with previous observations (Noble, Wei, et al. 2018), with an average of 86, 99 and 80% of particles associated with the AWI across the apoferritin, HSPD1 and ribosome data, respectively (Figure 2, Table S1 & Figure S1).

**Figure 2.**
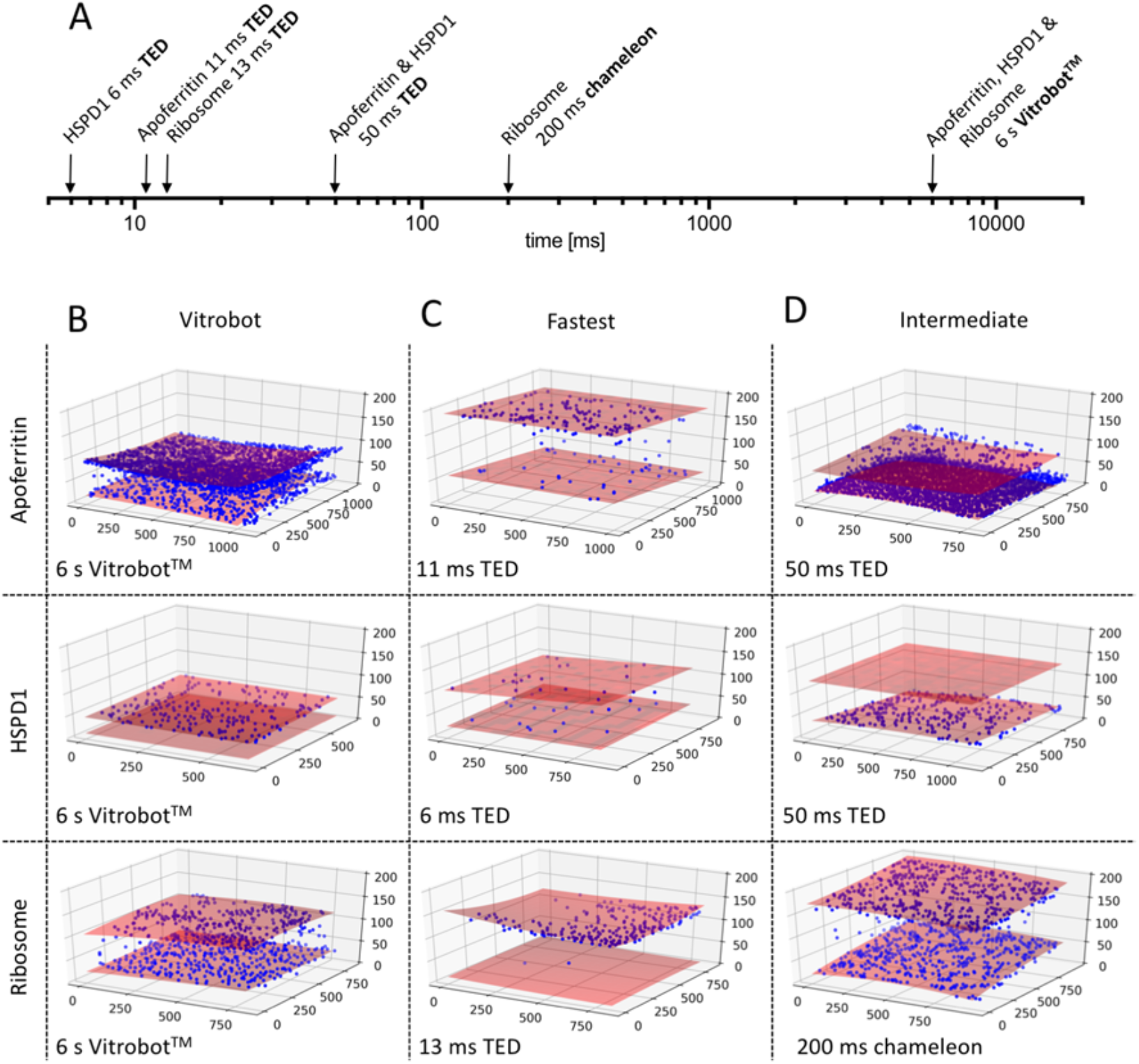
Visualisation of particle partitioning at AWI using cryoET. (A) Timescale of grid preparation for tomography samples. Representative tomograms of apoferritin, HSPD1 and ribosome grids prepared at standard blotting speed (Vitrobot™, time given is Vitrobot™ ‘blot time’) (B), fastest time points (TED) (C) and intermediate time points (D). Red shaded area indicates AWI, and blue spheres particle location. The axis indicates coordinates of the particle location in ice in nm. Time scale from sample preparation to vitrification and sample preparation device shown in the bottom of the box. Full raw data can be seen in Figure 2, Supplement 1.

To investigate trends in particle distribution on different timescales of vitrification, the TED was used to vitrify grids on ‘fast’ timescales (6-13 ms), and the TED and chameleon were used to vitrify grids on ‘intermediate’ timescales (50-200 ms). The majority of the particles partitioned to the AWI on the TED ‘fast’ timescale (apoferritin: 75% at 11 ms, HSPD1: 89% at 6 ms, and ribosome: 96% at 13 ms), although it should be noted the TED data showed a greater variability compared with Vitrobot™ data (Figure 2, Figure. S1). On the ‘intermediate’ timescale, TED grids of apoferritin (50 ms) and HSPD1 (50 ms) displayed large variability across different tomograms of the same specimen, however the majority of particles interacted with the AWI (67% and 95% for apoferritin and HSPD1, respectively). ‘Intermediate’ timescale chameleon grids of ribosome (200 ms) displayed 94% of sample interacting with the AWI.

The ‘fast’ TED data demonstrate that even on the fastest timescales we could investigate using this device and in thick ice (up to ~180 nm), the interaction with the AWI is not eliminated. This is perhaps unsurprising given that calculations suggest particles will interact 10-100 times with the AWI within 1 ms and for some proteins this interaction results in sequestering at the AWI (Naydenova & Russo 2017). It should be noted that TED generally produces thicker ice, especially at faster dispense-to-plunge times, as the TED relies on droplet spreading upon contact with the grid to produce areas sufficiently thin to image (Table S1). For the apoferritin grids prepared using the TED, we observed interesting trends in surface protein aggregates at 11 ms comparing to 50 ms. At 11 ms, small aggregates of ~10-50 particles were observed, which appeared to be much larger at 50 ms where they consisted of ~100s of particles. Protein aggregates were only observed at the AWI (Figure. S2).

When considering the spraying devices across various timescales and Vitrobot™ blotting data together, the trend of a reduction in particles at the AWI at faster freezing times holds true for the apoferritin and HSPD1 samples (Figure S1), although more variability is seen in the intermediate timepoints of grids made on the TED. Interestingly, the ribosome data show the opposite trend, with increased partitioning to the AWI at 13 ms compared to the blotted grid. A general observation across all sample preparation techniques (TED, chameleon, Vitrobot™) was the presence of asymmetry in particle distribution in some tomograms, i.e. one AWI face was highly populated while the other was not (Figure 2), as previously reported for the Vitrobot™ and Spotiton (Noble, Dandey, et al. 2018).

### Concentration of particles

Previous studies suggested that there is a variation in the concentration of the necessary amount of sample required to achieve similar particle numbers in frozen grids when using the Vitrobot™, TED and chameleon. We used tomograms to calculate particle concentration within the thin film on frozen grids, which was compared to the concentration of the protein solution used during grid preparation using the Vitrobot™, TED, and chameleon.

For Vitrobot™ blotted grids, there was a large increase, or concentrating effect, with average 3, 21 and 24-fold increases in particle numbers for apoferritin, HSPD1 and ribosomes, respectively (Figure 3). This interesting result demonstrates a previously unreported advantageous sample concentration effect of blotting methods. To interrogate this further, a comparison was made between a theoretical model thin film and the observed data. A model thin film was generated by placing particles representing the actual concentration of sample applied to the grid with randomly generated coordinates (within the confines of the thin film, assuming no concentration change and no affinity for the AWI) (Figure S3). For apoferritin, the model data matched remarkably well the experimental data at distances away from the AWI (> 10 nm), indicating that the concentration effect seen in Vitrobot™ blotted grids of apoferritin exclusively stems from particles bound to the AWI.

**Figure 3.**
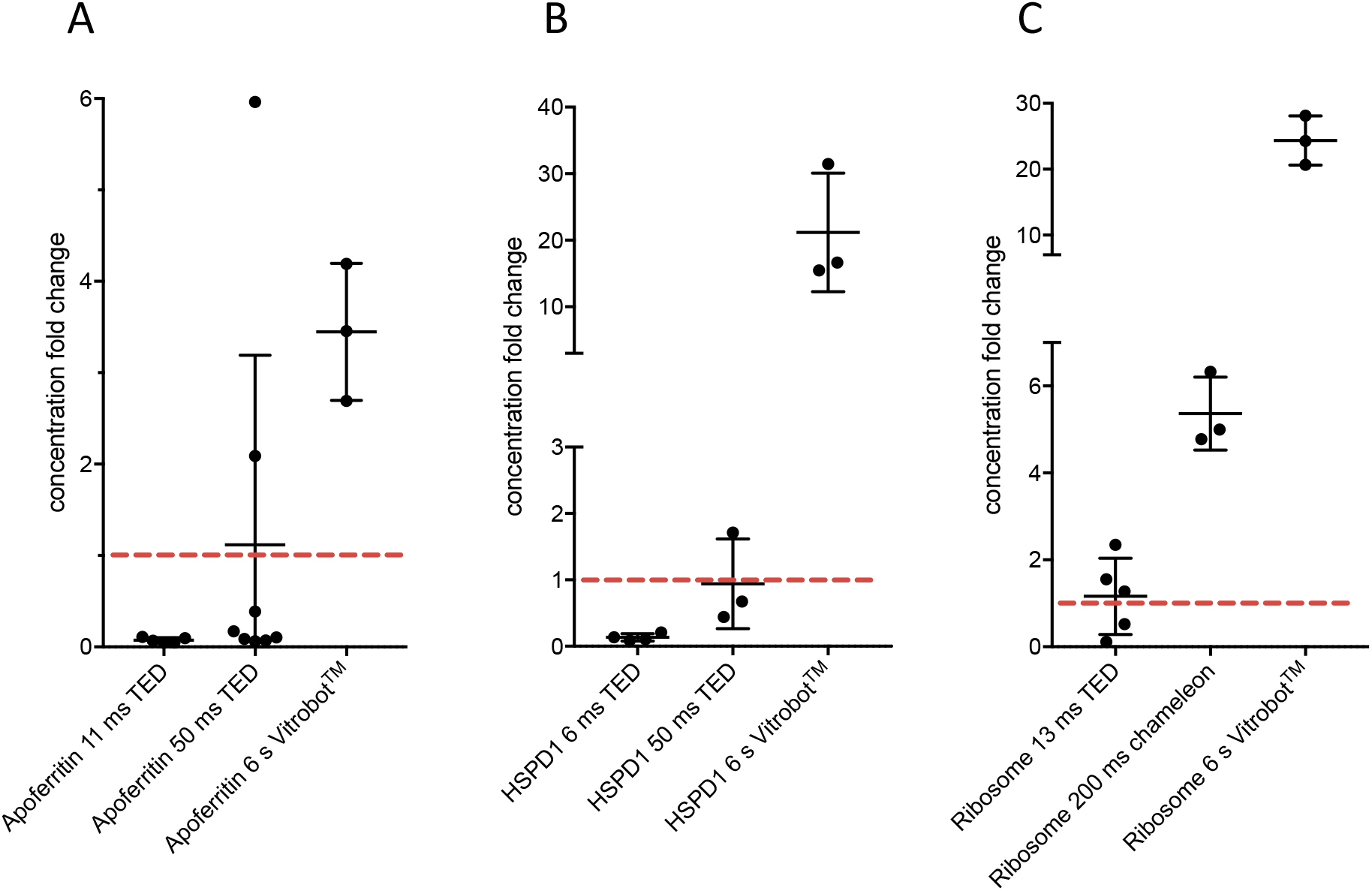
Apparent change in protein concentrations in the thin film at varying timepoints and vitrification devices. Particle concentrations in thin film as determined from tomograms for apoferritin (A), HSPD1 (B) and Ribosomes (C). Vitrification device and timescales identified on graph. Red line indicates the concentration in solution (applied concentration).

For the TED, we hypothesised that there should be no particle concentration or dilution effects, as the droplets land on the grid without liquid being drawn away as in the case of both the Vitrobot™ (filter paper) and chameleon (self-wicking grids). Using the TED, at 50 ms for HSPD1 and apoferritin (and 13 ms for ribosome), we do indeed see, on average, the number of particles we would expect given the concentration of protein applied. This indicates that there are no significant concentrating effects for TED at these timepoints. However, there is large variability in the 50 ms apoferritin data compared to the 50 ms HSPD1 data. Interestingly, the ‘fast’ apoferritin and HSPD1 data both show a large depletion of particles (14 and 7-fold respectively). Data from the chameleon on the ribosome sample at 200 ms shows a substantial concentrating effect (5-fold), but much reduced compared to the Vitrobot™ data.

### Orientation and angular distribution of HSPD1

HSPD1 is known to adopt strong preferred orientation when prepared using standard blot-freezing methods. We examined HSPD1 angular orientation using the TED at 6 and 50 ms, the chameleon at 54 ms and the Vitrobot™ (Figure 4). Single particle datasets for each timepoint and device were collected and combined after pre-processing. 2D- and 3D-classifications were performed on the combined data to impose the same class selection criteria on all datasets and the consensus structure was determined. From this, the angular assignments for particles that were frozen using each device at the specific timepoints, were extracted to analyse trends in preferred orientation (Figure 4C, Figure S4).

**Figure 4.**
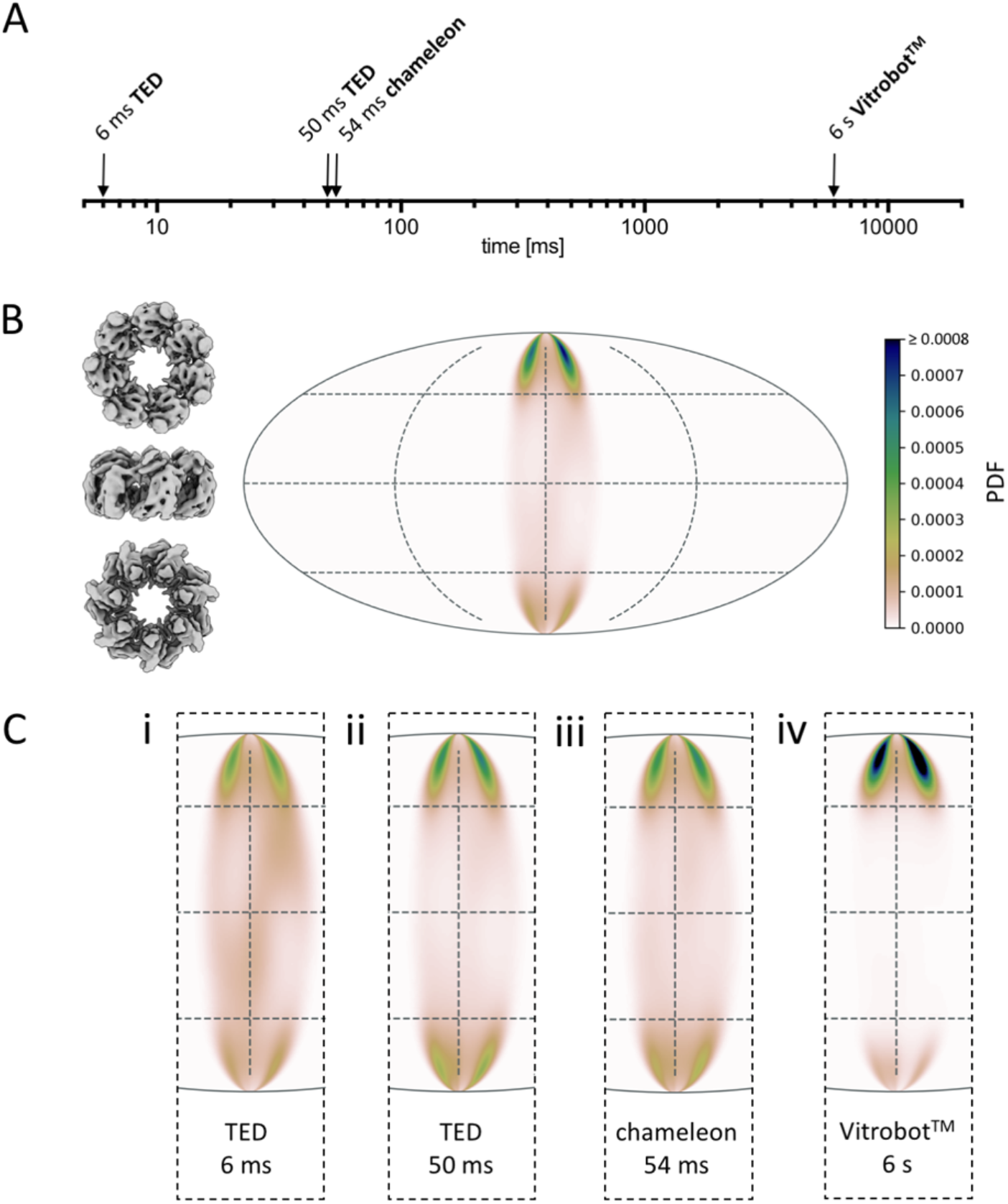
Angular orientation of HSPD1 over varying timepoints and vitrification devices. (A) Timescale of grid preparation for HSPD1 samples analysed for angular distribution. (B) Preferred orientation of HSPD1 of the combined data, showing an angular orientation distribution map (Mollweide projection) of the C7 symmetric reconstruction so only 1/7 of the area is occupied. Views of HSPD1 on the left show the approximate corresponding orientation, with these data dominated by the top view. (C) Orientation distribution maps for HSPD1 data collected from samples prepared with TED 6ms (i), TED 50 ms (ii), chameleon 54 ms (iii) and Vitrobot™ 6 seconds (iv). The normalised probability density function (PDF) approximates the probability to find a particle in a certain orientation. The colour scale is the same in (B) and (C).

As expected for HSPD1, strong preferred orientation was seen, with the ‘top’ and ‘bottom’ projections dominating the particle views present in all data collected. The quality of the consensus 3D reconstruction suffered from the anisotropy of views, as seen in the Z-directional FSC (Figure S5). The Vitrobot™ blotted sample (Figure 4C) showed the strongest preferred orientation. By increasing the speed of grid making using either the chameleon or TED, more angular distributions were available compared to the standard blotted grid. Reducing the time-delay further, from 50 ms to 6 ms on the TED, provided further minor improvements in angular distribution, although the data were still dominated by preferred views.

Due to variations between datasets, such as ice thickness and particle number, it is not possible to draw comparisons between the freezing devices used and resolution outcomes. Instead we limit comparisons to the range of angular distributions. For example, the reconstruction from the 6 ms TED data, which had a greater angular distribution, is limited in resolution to approximately 7 Å. This is likely due to increased ice thickness compared to the other datasets (Table S1); other reconstructions are likely resolution-limited due to low particle numbers or ice thickness (Figure S4).

### Orientation and angular distribution of ribosomes

A sample containing the 30S, 50S and 70S ribosomes was used to investigate the angular distributions of three related specimens in one dataset to keep as many parameters constant as possible (ice quality etc.). Applying the same approach used to examine HSPD1 angular distribution, we collected single particle datasets for ribosome samples prepared with TED (13 ms), chameleon (54 and 200 ms), and Vitrobot™ blotted samples (Figure 5, Figure S6).

**Figure 5.**
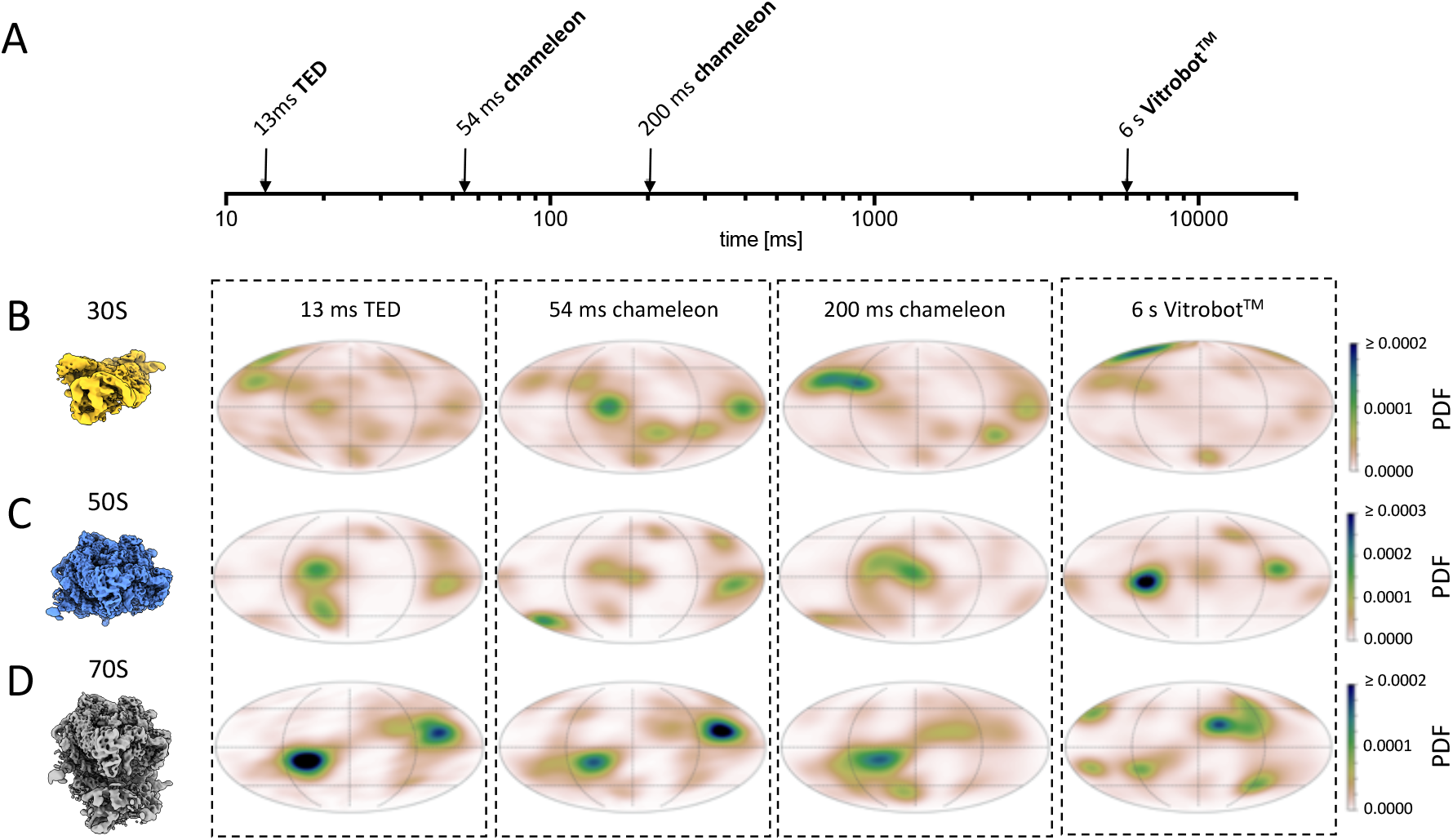
Ribosome angular orientation over varying timepoints and vitrification devices. (A) Timescale of grid preparation for ribosome angular SPA samples. Orientation distribution maps for 30S (B) 50S (C) and 70S (D) samples prepared using stated vitrification device and timescale. As in Figure 4, showing the normalised probability density function (PDF) in Mollweide projection to approximate the probability to find a particle in a certain orientation.

The 30S subunit showed a clear correlation between speed of grid preparation and improved angular distribution (Figure 5B). This trend was also present in the 50S subunit data, although it is not as pronounced (Figure 5C). Interestingly, this trend is not present for the full ribosome; instead the greatest angular distribution was observed from grids prepared using the Vitrobot™ (Figure 5D). Taking the datasets through the processing pipeline, none of the ribosome reconstructions appear to be limited in resolution by angular orientations, and the trends observed in resolution for each of the sample preparation times and methods appear to link most closely to the particle number (Figure S6 & S7).

Confirming AWI interactions can induce complex dissociation, we observe a number of ribosomal subunits which are resolved at early, but not later timepoints. Density for the 50 S ribosomal protein L31 is lost in the 70S and 50S ribosome structures in a time-dependent manner (Figure 6B & D). In the grids made in ≤ 54 ms, using both TED and chameleon, the L31 subunit is clearly present. However, in those grids made at 200 ms and 6 s, the L31 subunit is absent within the EM maps when viewed at the same and lower threshold as the fast plunge structures (Figure 6). The 30S and 70S reconstructions show that 30S ribosomal protein S2 also dissociates in a time-dependent manner. Interestingly, ribosomal protein S2 persists for a longer timeframe than the L31 subunit, only disappearing in the Vitrobot™ prepared grids while present in the TED and chameleon datasets (Figure 6F). The density for 50S ribosomal protein L9 behaves in a similar fashion, the difference is more pronounced in the 70S reconstructions, it is present at ≤ 200 ms but missing in the 6 s reconstruction.

**Figure 6.**
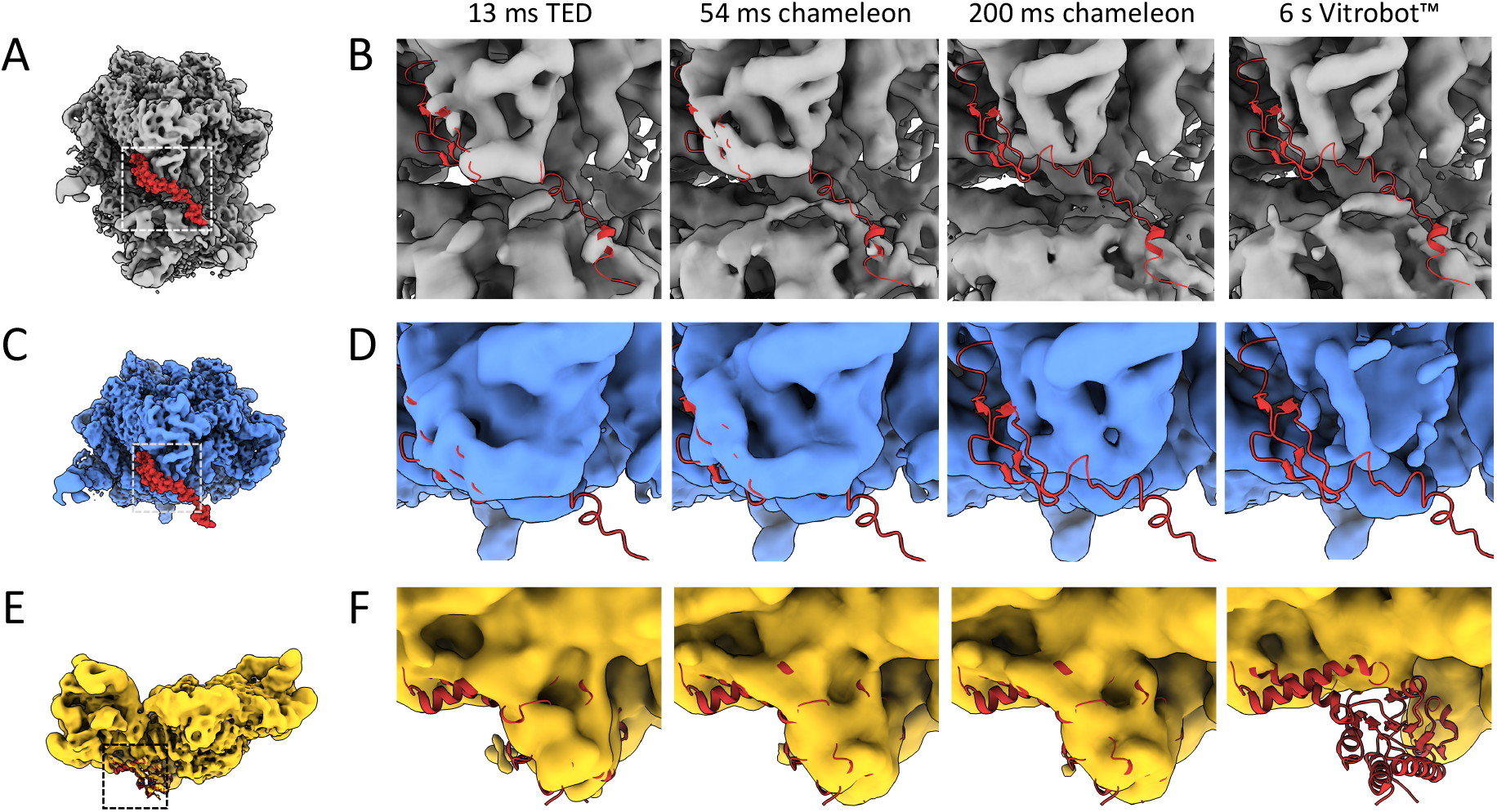
Dissociation of ribosomal subunits over varying timepoints and vitrification devices. (A) Position of ribosomal protein L31 (from PDB 6OSK) in the 70S ribosome. (B) Density for L31 in 70S reconstructions compared between all timepoints. While present in ‘fast’ reconstructions, the density is absent in the 200ms or Vitrobot™ reconstructions. (C) Position of ribosomal protein L31 (from PDB 6OSK) in the 50S ribosome. (D) Density for L31 in 50S reconstructions compared between all timepoints showing the same trend as in (B). (E) Position of ribosomal protein S2 in the 30S subunit (from PDB 6O7K). (F) Density for S2 in 30S reconstructions compared between compared between all timepoints. The S2 density is missing in the Vitrobot™, but present in all other reconstructions. 70S in grey, 50S in blue and 30S in yellow, all maps in B, D and F are shown at threshold of 3σ.

## Discussion

### AWI partitioning

The physics of diffusion and AWI interactions cannot be outrun using technology currently available (to the best of our knowledge) for cryoEM sample preparation. Even in the fastest cases of grid vitrification in our study (6 ms) and using different approaches (blotting vs spraying), the majority of particles still partitioned to the AWI. Considering AWI partitioning data from the three specimens we examined, apoferritin, HSPD1 and ribosomes, conflicting lessons can be learnt from each. HSPD1 data suggest that the faster grids are prepared, the fewer particles partition to the AWI (Figure 2, Figure S1). The ribosome data suggest the precise opposite; the faster the grids are prepared, the more particles partition to the AWI (Figure 2, Figure S1). The apoferritin data are the most variable and provide the least clear picture across different timescales, which may be partially explained by the propensity of apoferritin to form ‘rafts’ at the AWI (discussed below in ‘changes in particle concentration’).

Overall, altering speed of grid preparation could be one mechanism to influence AWI partitioning, but the effects of this are not linear and are difficult to predict across different specimens. A greater understanding of the factors that may influence partitioning, including specimen polarity, stability, and buffer composition, along with more information about how different specimens respond to the thin film environment over time, may enable better predictions of specimen behaviour prior to freezing in the future.

### Changes in angular distribution

Even though most particles could not be prevented from locating at the AWI, small (HSPD1) to very large (30S, 50S ribosome) changes in the angular distribution of particle over 10s to 100s of milliseconds were observed (Figure 4, Figure 5). One of the most fascinating aspects of these data is the time-related change in angular distribution related to timescale, and how this varies depending on the sample. These data suggest that for a given specimen, there may be a fast (<10 ms) stage where the protein initially partitions to the AWI, followed by a slower stage where the particle explores its energy landscape before settling into a local energy minimum. For some specimens, there may be a distinct orientation (leading to preferred orientation), and for other specimens it may be a variety of orientations (Figure 7). The timescale in this second, slower stage is likely to vary from specimen to specimen.

**Figure 7.**
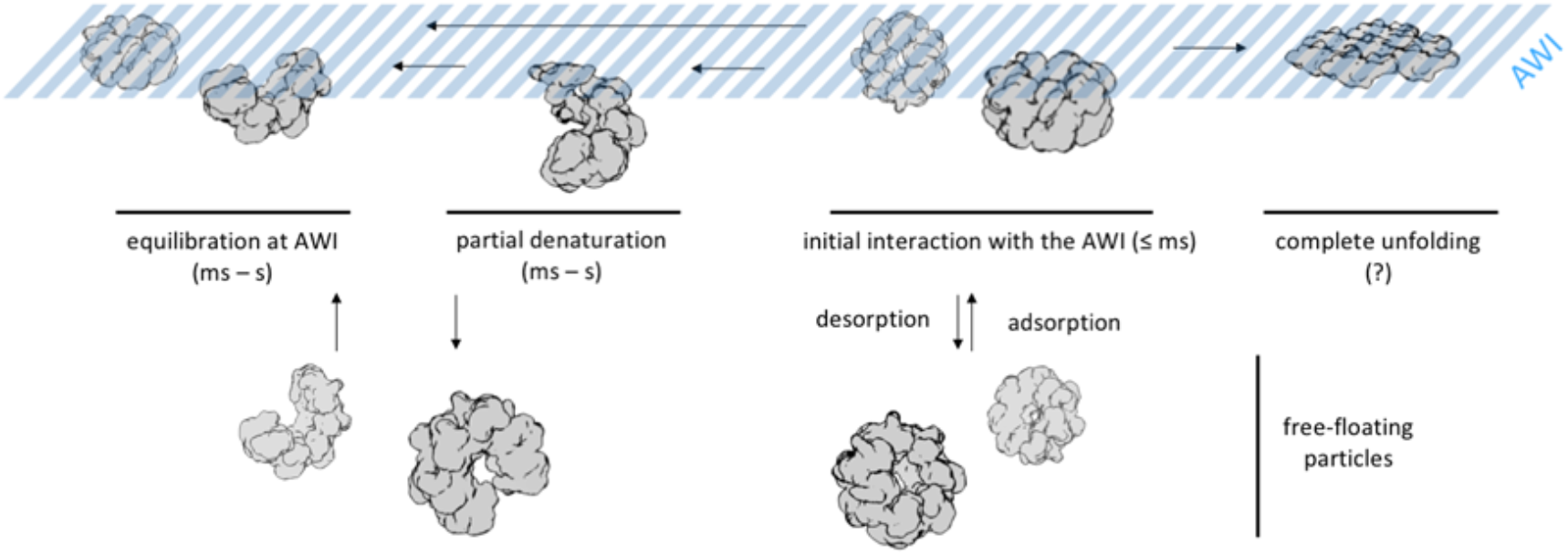
Proposed model of protein-AWI interactions. The initial interaction with the AWI (adsorption) is fast. equilibration of the protein-AWI system, however, is slower as it involves processes like desorption, partial, or complete denaturation which occur on various timescales (ms – s) and are thought to be highly protein-dependent (desorption from the AWI, rate of unfolding).

### Particle damage over time

Another factor that must be considered at the AWI, is the partial or full denaturation of protein specimens (D’Imprima et al. 2019). The timeframe of such denaturation at the AWI is likely to depend on many factors. The ribosome data suggests that denaturation during sample preparation can occur on the 100s-1000s of milliseconds timescale. In 70S and 50S ribosomes, the L31 subunit is only present at timepoints < 200 ms, while 30S ribosomal subunit S2 and 50S subunit L9 are still present at 200 ms (Figure 6). In agreement with our data, a recent study has shown 30S ribosomal subunit S2 is present when grids have carbon support, but dissociated on unsupported grids (Jahagirdar, D. et al., 2020). These data suggest that the timescale of partial denaturation at the AWI is highly specimen-dependent, and extends into the timeframe accessible by various grid preparation methods. This timescale suggests that partial denaturation may be an effect of the slower energy landscape exploration, helping to explain how particle orientation can change over longer timescales than partitioning to the AWI takes (Figure 7).

### Concentrating effect of Vitrobot™ blotting

One of the most striking results was the change in concentration due to blotting as compared with spraying (Figure 3). These data clearly demonstrate that for the specimens we have examined, the Vitrobot™ blotting approach greatly enriches the thin film with particles, for the specimens examined here. And indeed, that the AWI may be responsible for the concentration of particles in the thin film, which in many systems is required to achieve a viable number of particles per micrograph. It should also be noted that the degree of concentration is sample-dependent. This may go some way towards explaining the experience of many cryoEM researchers in ascertaining the ‘right’ concentration of protein to use for their system. Adsorption to the grid support may also have a significant impact on apparent particle concentration in the imageable areas which requires further investigation.

### Changes in particle concentration due to speed of grid preparation

Across both TED and chameleon, higher concentrations of specimen were necessary at faster timepoints. However, specifically with TED at the ‘fast’ timescales, a depletion of particles for apoferritin and HSPD1 was observed (Figure 3). The apoferritin data from TED display greater variability relative to the other samples, which could be linked to the formation of surface aggregates that were also observed in these data (Figure S2). Surface aggregates, or particle ‘rafts’, may begin to form while the droplet is traveling from nozzle to grid in TED (~0.5 ms). The size of the aggregates may be time-dependent with increases in size occurring at longer timescales. These rafts create locally high concentrations of particles on a single interface (Figure S2 Ci & Cii). Occasionally, large rafts are found in the thin areas chosen for data collection, and are likely also present in the thick regions unsuitable for imaging by TEM. This rafting behaviour may explain why, on average, the expected number of particles are present in TED 50 ms apoferritin samples, but with large variability in concentration from area to area. A small number of examples of the ‘raft’ effect were also observed for the 13 ms ribosome grids, but not for HSPD1 data indicating the presence of rafts is sample-dependent, while its severity is time dependent. No ‘rafting’ was seen in grids prepared using chameleon or the Vitrobot™.

A major unexplained aspect of these data is that for HSPD1 and apoferritin at the ‘fast’ TED time points (11 and 6 ms respectively), a large depletion in the concentration of particles compared with slower speeds was observed (Figure 3). We propose the following two hypotheses to explain these observations:

1. The first hypothesis relates to the variability of droplet size, a feature specific to the TED (Figure 1B). The droplets have variable surface-to-volume ratios, so in smaller droplets particles would be more likely to interact with the AWI from the moment the droplet is formed and travels to the grid. If denaturation occurs at this interface, then apparent protein concentration would decrease in this droplet and become lower than what would have been observed in larger droplets where the surface to volume ratio would favour proportionally fewer AWI interactions. Once on the grid and frozen, smaller droplets are more likely to be imaged, especially at ‘fast’ grid preparation speeds, as they are more likely to result in thin ice. The larger droplets, which, according to this hypothesis, would contain closer to the expected number of particles, cannot be imaged at these ‘fast’ timepoints as they will result in ice that is too thick. At intermediate grid preparation timescales, the smaller droplets may have disappeared (due to film thinning), while the larger droplets have thinned to suitable thicknesses. This would explain the apparent depletion of particles observed at faster speeds, and the ‘reappearance’ of particles at intermediate timepoints.
2. Particle denaturation over time may also contribute to the observed concentration differences. On TED prepared grids, each frozen droplet has a thick central part that subsequently spreads out into a thin layer at the edge of the droplet. The apparent depletion of particles at the ‘fast’ timepoints may be due to the loss of proteins in the formation of a sacrificial denatured protein layer. Particles could then diffuse from thicker areas in the droplet to repopulate thin, imageable areas, causing the ‘reappearance’ of particles at intermediate timepoints. Theoretical calculations suggest that protein denaturation may occur on a sub-millisecond timescale (Raffaini & Ganazzoli 2010) in addition to the longer timescales for partial denaturation seen here (Figure 7).

Formation of a sacrificial layer of denatured protein has been shown for apoferritin (Yoshimura, Hideyuki, et al. 1994), but the timescale of complete particle denaturation on cryoEM grids remains an open question. There may be alternative explanations for these data, and the hypotheses presented are not mutually exclusive. It is likely that there is an interplay between multiple mechanisms on a specimen-dependent basis. It is only with additional information on these trends across many specimens, added to these initial data that a better understanding of particle behaviour in thin films can be achieved.

In conclusion these data go some way to offering an explanation to those cryoEM researchers who have experienced huge variability in cryoEM sample preparation between biological specimens. General trends indicate speed may ameliorate some of the adverse effects of the AWI, thus providing a significant improvement in intact or non-preferentially oriented particles. However, this speed may come at the price of a higher required sample concentration, with data suggesting that, the faster grids are prepared, the higher the concentration of protein required. This effect may seem exacerbated given the concentrating effect currently enjoyed when using Vitrobot™ blotting to prepare samples.

While much is still unknown about the behaviour of particle in thin films, a general model can be used to summarise the aforementioned ideas (Figure 7). First, diffusion dictates the rate at which a particle interacts with the AWI. This is an initial fast phase, occurring within ≤ 1 ms of the thin film forming. Each specimen will then have its own on-off rate and local energy minima at the AWI, determining how likely it is for the protein to disassociate back into bulk solution. Next, negative aspects of the AWI may take place with partial denaturation or dissociation of parts of the molecule, and/or adoption of preferred orientations. However, the timescales of this final equilibrium will likely be highly specimen specific. This model explains why no ‘silver bullet’ has yet been developed to generically tackle cryoEM sample preparation for every specimen. Speed of grid preparation, grid types, use of surfactants, continuous and engineered supports and new grid making technologies will all have a role to play as the field moves towards the development of generically useful approaches for cryoEM sample preparation, but in the meantime, they can be used as individual tools along the research path toward an optimised cryoEM grid.

## Materials and Methods

### Sample preparation

Horse spleen apoferritin was purchased from Sigma Aldrich (A3660), and exchanged into 30 mM HEPES, 150 mM NaCl pH 7.5 by ultrafiltration using 100 kDa molecular weight cut-off (MWCO) spin concentrator tubes (Vivaspin, Sartorius). Protein concentration was then determined using absorbance at 280 nm (ε_280_ = 14,565 mol^−1^cm^−1^, MW= 18.5 kDa and homo-24-mer stoichiometry). For grid preparation, apoferritin was diluted to the target concentration in 30 mM HEPES, 150 mM NaCl pH 7.5.

*E. coli* ribosome sample was purchased from New England Biolabs (P0763S), provided at a stock concentration of 33 mg/mL (= approx. 25 μM assuming an average molecular weight of 1.34 MDa (Van Holde & Hill 1974). For grid preparation, ribosomes were diluted to the target concentration using 50 mM HEPES, 8 mM MgAc_2_, 100 mM KAc pH 7.5.

Mature human mitochondrial heat shock protein family D member 1 (HSPD1) was expressed in *E. coli* BL21 DE3 and purified based on a modified version previously described protocol (Viitanen et al. 1998). The expression plasmid was kindly provided by Dr Hao Shao and Dr Jason Gestwicki (UCSF). Competent *E. coli* BL21 DE3 were transformed with the plasmid using the heat-shock method. Two flasks of 1 L TB media were inoculated with 2 × 20 mL overnight culture, incubated for 2.5 h at 37 °C, 200 rpm until OD_600_ reached 0.8. Expression was induced by adding 250 μM IPTG and cells were further incubated for 4 h at 37 °C, 180 rpm, then cells were harvested by centrifugation (10 min, 4000 rpm) and stored at −80 °C.

All purification steps until reconstitution were done on ice or at 4 °C. The cell pellet was thawed on ice and resuspended in 40 mL lysis buffer (50 mM Tris, 500 mM NaCl, 10 mM imidazole pH 8), supplemented with 1 mM PMSF and protease inhibitor cocktail (set V, Calbiochem). The cells were further resuspended with 4 strokes in a dounce homogeniser and lysed with a sonicator (35% amplitude, 30 sec on/off, 10 min total). Cell debris was pelleted by centrifugation (30 min, 17,000 rpm) and the supernatant applied to 7 mL packed, equilibrated Ni-NTA resin. The protein-bound resin was washed with 200 mL lysis buffer and 200 mL wash buffer (50 mM Tris, 300 mM NaCl, 50 mM imidazole pH 8). The protein was eluted with 20 mL eluting buffer (50 mM Tris, 300 mM NaCl, 300 mM imidazole pH 8). To remove the His6-tag, DTT (final concentration 1 mM) and TEV protease (1.6 mg per 10 mL, 3.2 mg in total) were added and the mixture was incubated for 4 h at room temperature. The cleavage products were dialysed against 4 L dialysis buffer (50 mM Tris, 150 mM NaCl pH 7.5) in a 10 kDa MWCO membrane overnight. TEV protease and His6-tag were removed by incubation with 3 mL of preequilibrated Ni-NTA resin for 1 h in lysis buffer. The flowthrough was collected, 10% (v/v) glycerol was added and the protein concentrated to 20 – 30 mg/mL in a 10 kDa MWCO spin concentrator (Vivaspin, Sartorius).

For reconstitution into its oligomeric form, 4 mL HSPD1 were mixed with 100 μL 1 M KCl, 100 μL 1 M Mg(OAc_2_) and 400 μL 50 mM Mg-ATP (pH 7). The reaction was incubated at 30 °C for 60-90 min. All following steps were done at room temperature. Precipitate was removed by centrifugation at 13,000 rpm for 10 min and the soluble fraction was loaded onto a HiLoad 16/600 Superdex 200 gel filtration column (GE Healthcare). Size exclusion chromatography was done in 50 mM Tris, 150 mM NaCl, 10 mM MgCl_2_ pH 7.7. The fractions corresponding to oligomeric HSPD1 (as determined by negative stain EM and SDS-PAGE) were collected, concentrated to 10-25 mg/mL with 10 kDa MWCO spin concentrators, supplemented with 5% (v/v) glycerol and frozen in liquid N_2_.

Protein concentrations of HSPD1 was determined by measuring absorbance at 280 nm (ε_280_ = 14,440 mol^−1^cm^−1^, MW = 58.2 kDa and homo-7mer stoichiometry). For grid preparation, HSPD1 was diluted to the target concentration in 50 mM Tris, 300 mM NaCl, 10 mM MgCl_2_ pH 8.

### Preparation of blotted grids

For specimens prepared by blotting, Quantifoil 300 mesh Cu R 1.2/1.3 holey carbon grids were glow-discharged in a Cressington 208 carbon coater with glow discharge unit at 10 mA and 0.1 mbar air pressure for 30 s. Grids were prepared using a Vitrobot™ mark IV (Thermo/FEI) with a blot force of 6 and a blot time of 6 s. The relative humidity (RH) was ≥ 90% and temperature 20 °C for ribosome and 4 °C for apoferritin and HSPD1. Concentrations for Vitrobot™ grid preparation were 20, 0.6 and 0.8 μM for apoferritin (24mer), HSPD1 (7mer) and ribosome, respectively. The applied sample volume was 3 μL for all blotted grids and the liquid ethane was used as cryogen in all cases.

### Fast preparation of grids using the TED

Fast grid preparation using the TED was done as previously described, using gas-dynamic virtual nozzles in spraying mode (Klebl, D.P. et al, 2020). Quantifoil 300 mesh Cu R 1.2/1.3 holey carbon grids were used after glow-discharge in a Cressington 208 carbon coater with glow discharge unit at 10 mA and 0.1 mbar air pressure for 99 s. In this TED setup, the droplets are small and fast and the delay between spray and deposition short (≤ 1 ms). The spray parameters were held approximately constant for all grids, using a liquid flowrate of 8.3 μL/s and an atomizer gas pressure between 1.5 and 2.0 bar. The nozzle design used was slightly different from the one previously described with the distance between liquid channel and nozzle outlet being 95 μm instead of 125 μm. PDMS sprayers were manufactured as previously described. Droplet speeds are high under these conditions (> 20 m/s) and the used nozzle-grid distance (during sample application) was low (7 - 10 mm). Therefore, to estimate exposure time of the thin film to the AWI, only the time between droplet impact on the grid and freezing was considered. Plunge speeds were measured using a linear potentiometer and the vertical distance between nozzle and liquid ethane surface was 1-3 cm and the plunge speed was ≤ 3 m/s. The humidity chamber was at ≥ 80 % RH and ambient temperature for grid preparation. Concentrations for TED grid preparation were 20, 11 and 2.5 μM for apoferritin (24mer), HSPD1 (7mer) and ribosome, respectively.

### Fast preparation of grids using the chameleon

For specimens prepared on the chameleon system, SPT Labtech 300 mesh Cu R 1.2/0.8 holey carbon self-wicking nanowire grids were used. Variable amounts of glow discharge in a Pelco Easiglow at 12 mA, 0.39 mbar air pressure were used to activate and control the wicking speed. Samples were held at 4 °C (apoferritin, ribosome) or 24 °C (HSPD1) until aspiration into the dispenser. Grids were prepared at a RH between 75% and 85% at ambient temperature. The applied sample volume for each stripe is ~6 nL. Concentrations for chameleon grid preparation were 5.5 and 2.5 μM for HSPD1 (7mer) and ribosome, respectively.

### Fiducial-less cryoET data collection and processing

All cryoET was collected in the Astbury Biostructure Laboratory in Leeds on Titan Krios II, using the Gatan K2 direct electron detector operated in counting mode and a Bioquantum energy filter. Data acquisition parameters are listed in Table S1.

Frames were motion-corrected with MotionCor2 (Zheng et al. 2017), stacked using an in-house script and tomograms were reconstructed using back projection in Imod after 4-fold binning to enhance the contrast (Mastronarde 1997; Kremer et al. 1996). Particles were manually picked using EMAN2 (Tang et al. 2007). Particle positions were then used to locate the AWI. In order to do this, the tomogram was divided into patches in the x/y-directions (4-16 patches depending on particle concentration). The particles at minimum and maximum Z-height were selected and used to fit a plane (first or second order polynomial, depending on the number of patches and visual inspection of the fit) which corresponds to the upper and lower AWI, respectively. Then, the closest distance was determined between either of the AWIs and each particle. Particles which were at a distance ≤ 10 nm to an AWI were classed as ‘bound’ to the AWI. For the majority of tomograms collected, the 10 nm threshold adequately allowed characterisation of the data, but for tomograms on areas of thick ice (> 80 nm)/where the AWI is not clearly defined, a threshold of 20 nm was more suitable. Ideal particle behaviour was modelled using the experimentally determined AWIs and randomly generating particle coordinates (number/volume corresponding to the respective concentration) in between the experimental ice layer. Then, distances between modelled particles and AWIs were determined.

### Single particle cryoEM data collection and processing

All single particle cryoEM data was collected in the Astbury Biostructure Laboratory in Leeds on Titan Krios I, equipped with a FEI Falcon III detector and operated in integrating mode. Data collection parameters are listed in Tables S3 and S3.

All single particle data processing was done in RELION 3 (Zivanov et al. 2018). Micrographs were corrected for beam-induced motion with MotionCor2 and the CTF was estimated using GCTF (Zhang 2016; Zheng et al. 2017). All further data processing was done as shown in Figure S4 and Figure S6 for HSPD1 and ribosome, respectively. Particles were picked using the general model in crYOLO (Wagner et al. 2019).

All HSPD1 datasets were combined after particle extraction (rescaled to 2.13 Å pixel size). One round each of 2D- and 3D-classification were used to clean the dataset. Consensus reconstructions with particles from all datasets were generated in C1 and C7 symmetry and used to determine angular distributions. Finally, the dataset was split into its original subsets and each subset of particles and used to generate a reconstruction using the assigned angles from the C7 consensus reconstruction.

Similarly, all ribosome datasets were combined after extraction and subjected to one round of 2D classification to remove ‘junk’ particles. Then, 3D classification was performed to separate the combined datasets into 70S, 50S and 30S subsets. Those subsets were cleaned up by an additional round of 2D classification (2 rounds for 30S) and a consensus reconstruction was generated including data from all 4 datasets for the three species (70S, 50S and 30S). The subset for each species was then further split into the original datasets, resulting in reconstructions for 70S, 50S and 30S for each timepoint.

The maps were visualised using ChimeraX (Goddard et al. 2018). Orientation distributions were visualized using a script adapted from Naydenova *et al.* (Naydenova & Russo 2017). The probability density function was estimated using kernel density estimation with a gaussian kernel at a fixed bandwidth of 10°, wider than the estimated angular accuracy in all cases (to avoid overinterpretation of angular distribution maps).

## Acknowledgements

We thank Dr Hao Shao and Dr Jason Gestwicki (UCSF) for providing the HSPD1 plasmid used in this work. DPK and MSCG are PhD students on the Wellcome Trust 4-year PhD programme in The Astbury Centre funded by The University of Leeds. DJW and DK were funded by the BBSRC (BB/P020208/1 & BB/P026397/1, respectively). The FEI Titan Krios microscopes were funded by the University of Leeds (UoL ABSL award) and Wellcome Trust (108466/Z/15/Z). Maps have been deposited within the EMDB; ribosome 30S at 13ms, 54ms, 200ms and blotted (10871, 10872, 10873 and 10874, respectively), ribosome 50S at 13ms, 54 ms, 200 ms and blotted (10875, 10876, 10877 and 10878, respectively), ribosome 70S at 13ms, 54 ms, 200ms and blotted (10879, 10880, 10881 and 10882) and HSPD1 at 6ms, 50ms, 54ms and blotted (10883, 10884, 10885 and 10886).

