## Supplemental figures and tables for "Need for speed: Examining protein behaviour during cryoEM grid preparation at different timescales"

**Fig. S1.**

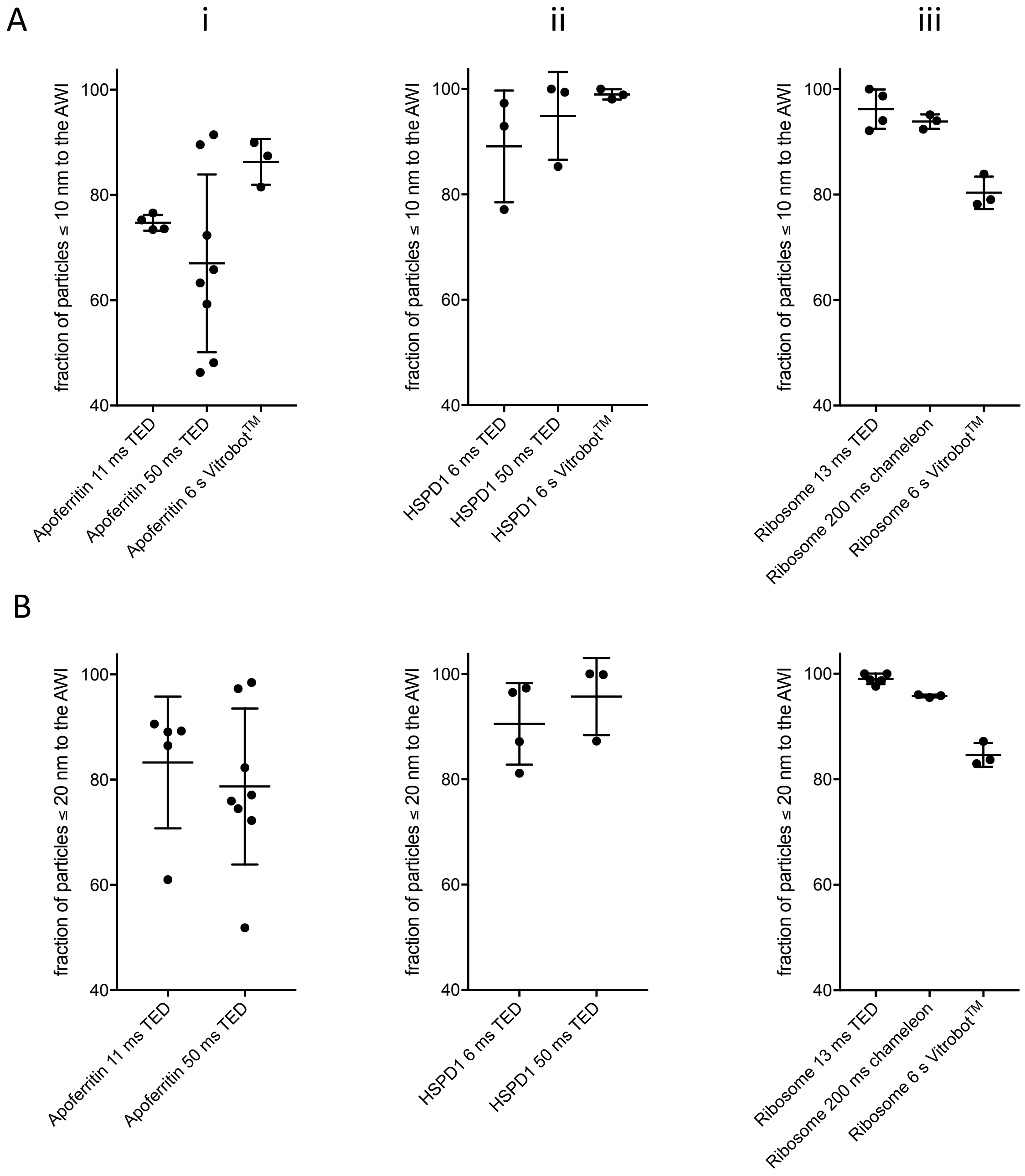


**Partitioning of particles to the AWI.** Fraction of particles partitioning to the AWI within 10 nm (A) or 20 nm (B) for apoferritin (i), HSPD1 (ii) and ribosomes (iii), with time and method of vitrification indicated. Some Vitrobot™ values were excluded from (B) because of low ice thickness. Shown are the individual data points, mean value and standard deviation.

**Fig. S2.**

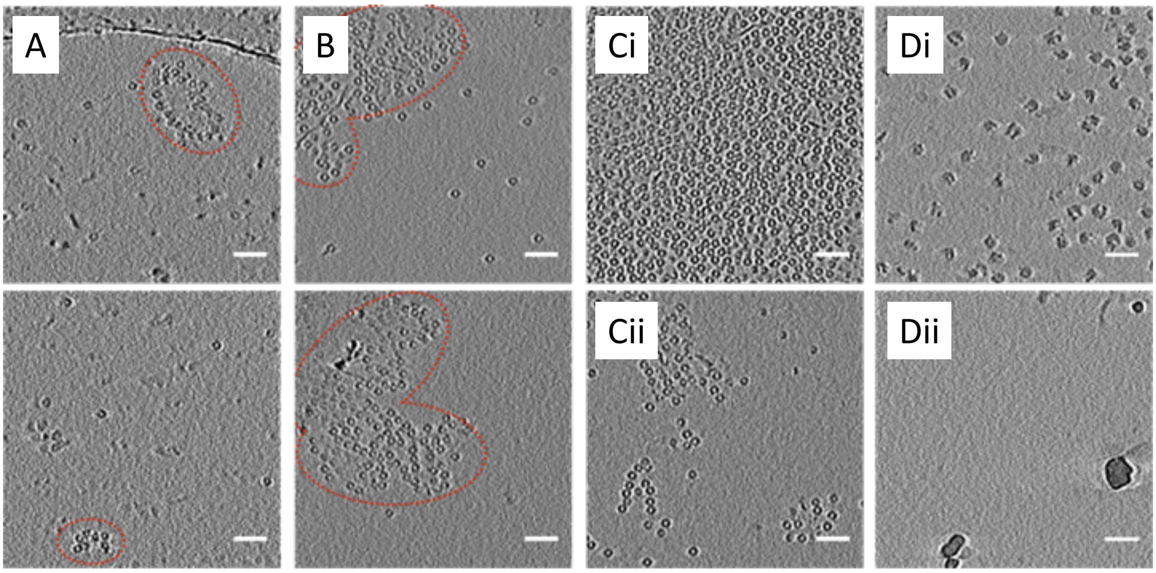


**Surface aggregates at varying timepoints.** Sections through reconstructed tomograms from TED grids, showing morphology of apoferritin aggregates at the AWI at (A) 11 ms or (B) 50 ms, outlined in red. Some tomograms showed highly asymmetric distributions of particles, with two interfaces from the same tomogram shown in (Ci) and (Cii) or (Di) and (Dii) for the apoferritin 50 ms and ribosome 13 ms sample, respectively. Scale bars 50 nm.

**Fig. S3.**

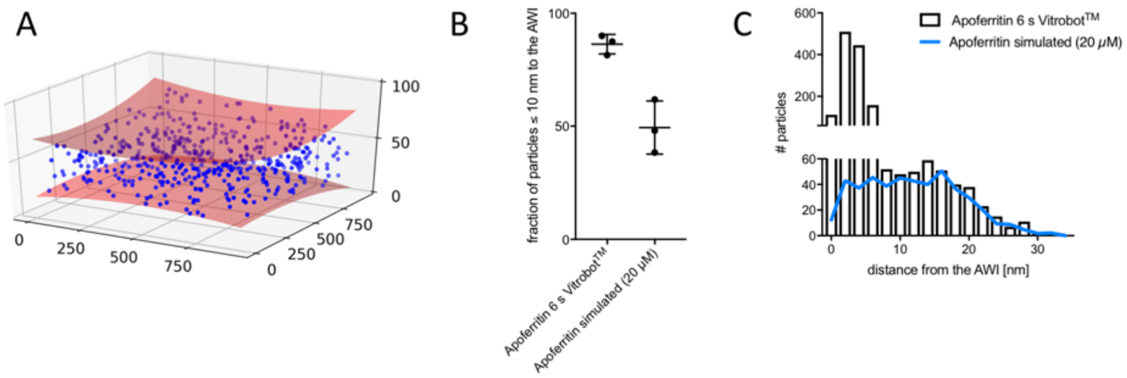


**Comparing modelled data to experimental apoferritin partitioning.** (A) Modelled distribution of particles (blue) in an ice layer (red), with coordinates indicated on axis (nm). AWIs were taken from an experimental tomogram and particle coordinates were randomly generated. This models a situation with no changes in concentration compared with sample applied, and no affinity for the AWI. (B) Comparison between modeled and experimental data for particle partitioning to the AWI. Particle concentration is 20 μM, $P = 0.007$. (C) Distributions of distances between particles and AWI for simulated (blue line) and experimental data (black bars) for apoferritin Vitrobot™ grids.

**Fig. S4.**

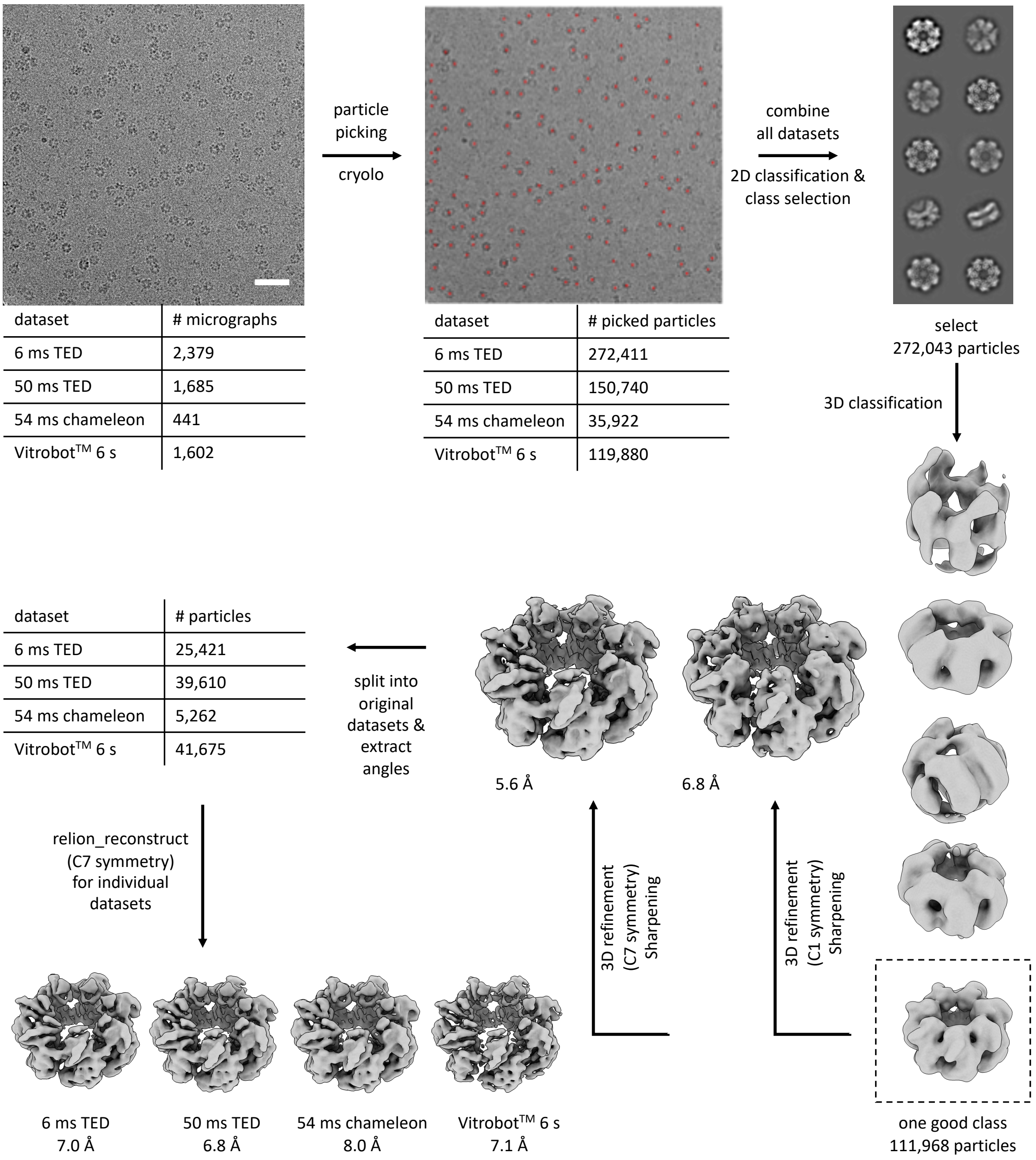


| dataset | # micrographs |
|---|---|
| 6 ms TED | 2,379 |
| 50 ms TED | 1,685 |
| 54 ms chameleon | 441 |
| Vitrobot™ 6 s | 1,602 |

| dataset | # picked particles |
|---|---|
| 6 ms TED | 272,411 |
| 50 ms TED | 150,740 |
| 54 ms chameleon | 35,922 |
| Vitrobot™ 6 s | 119,880 |

| dataset | # particles |
|---|---|
| 6 ms TED | 25,421 |
| 50 ms TED | 39,610 |
| 54 ms chameleon | 5,262 |
| Vitrobot™ 6 s | 41,675 |

**CryoEM image processing of HSPD1 data to yield angular distribution data.** Data processing pipelines for HSPD1 data showing particle numbers and corresponding 3D density maps and resolution for each sample preparation device and timescale analysed. Datasets have varying ice thicknesses, particle number and angular orientation and so resolutions cannot be directly compared.

**Fig. S5.**

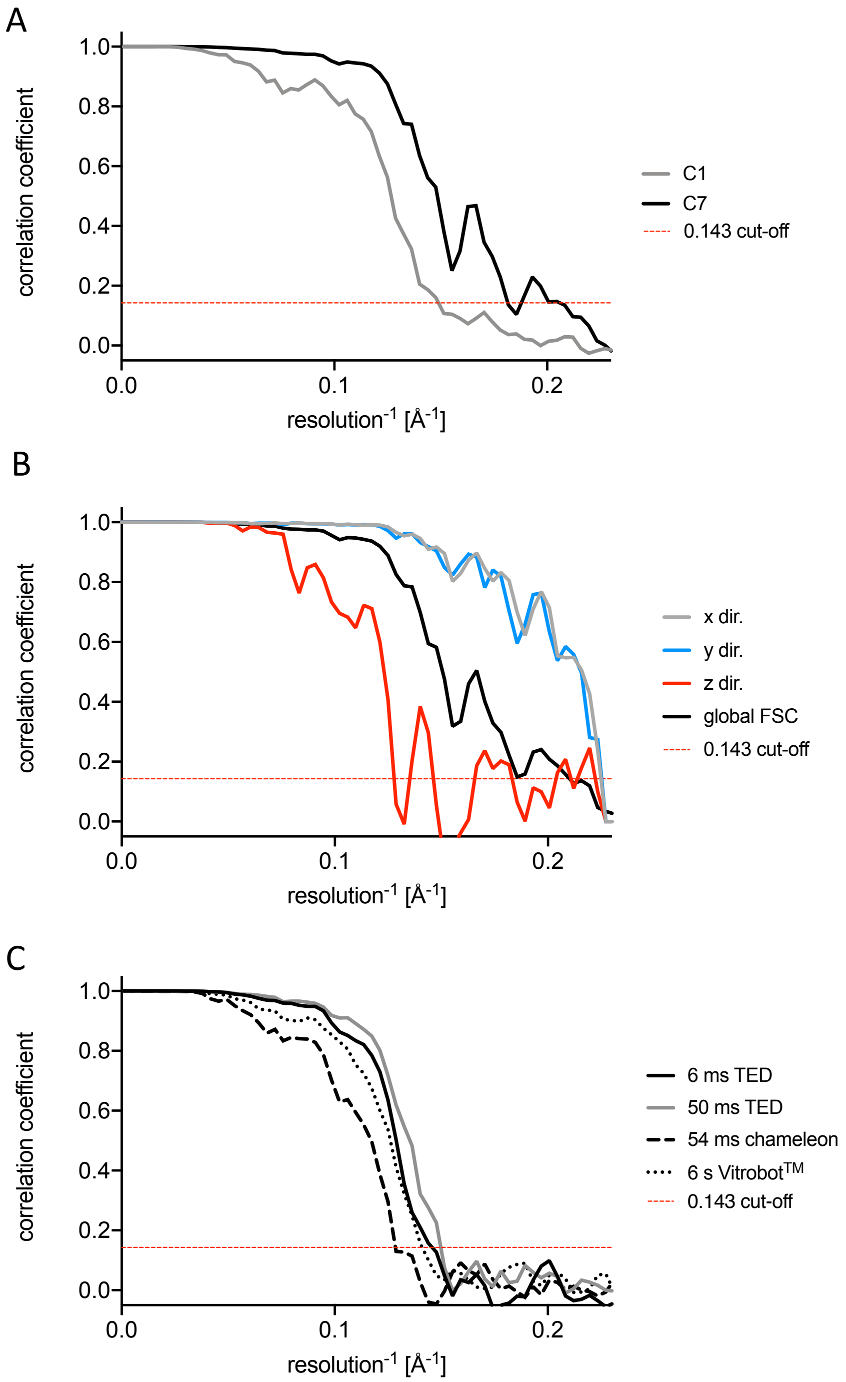


**FSC curves for HSPD1 consensus structures.** (A) FSC curves for masked maps of the consensus structure (all datasets combined) with and without symmetry. (B) 3D-FSC analysis of the same reconstruction, showing that resolution in the z-direction is limited through the lack of side views. Note that pixel size is 2.13 Å. (C) FSC curves for reconstructions for each individual dataset.

**Fig. S6.**

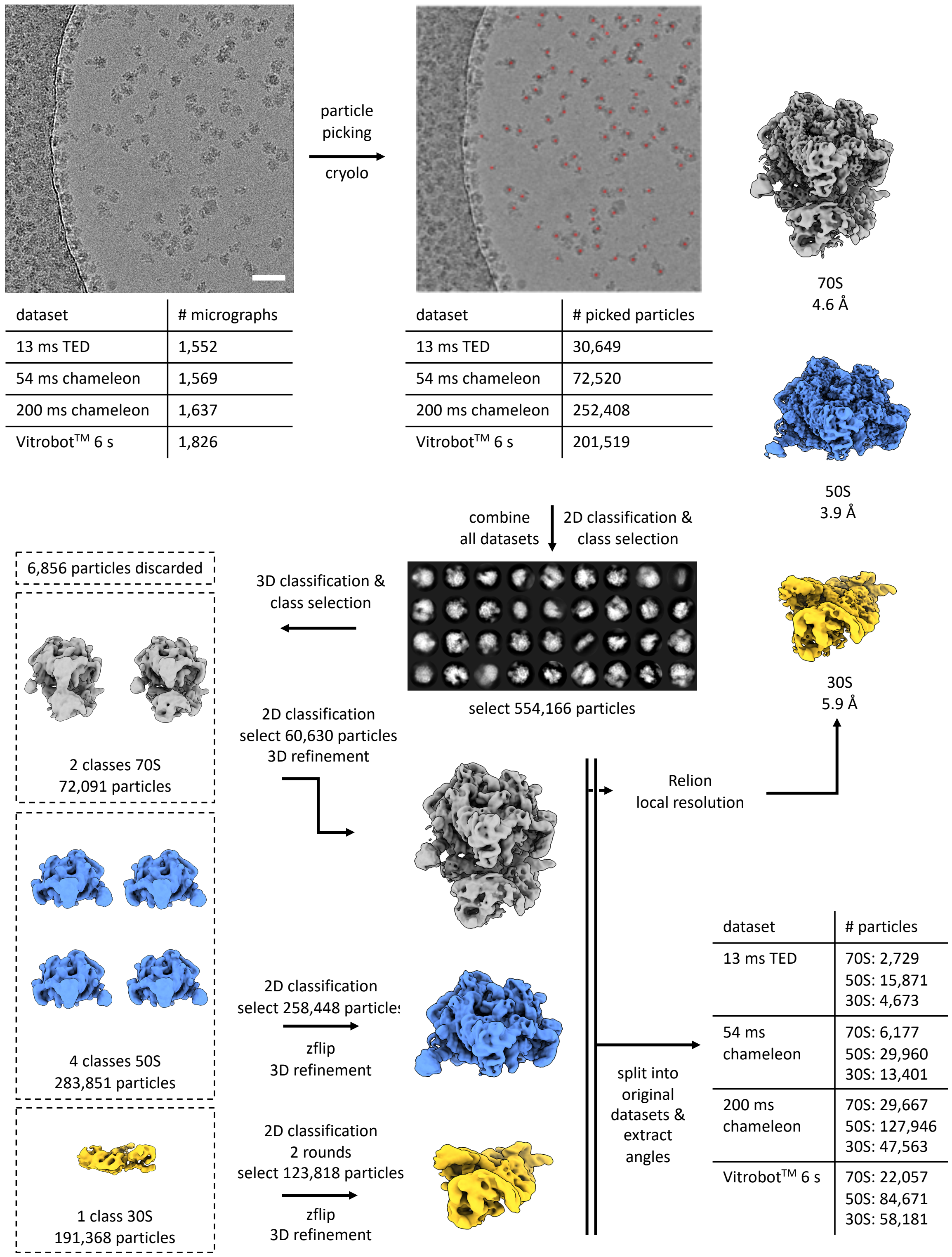


**CryoEM image processing of ribosome data to yield angular distribution data.** Data processing pipelines for ribosome angular orientation maps. Micrographs shown are representative of the type of micrograph used.

**Fig. S7.**

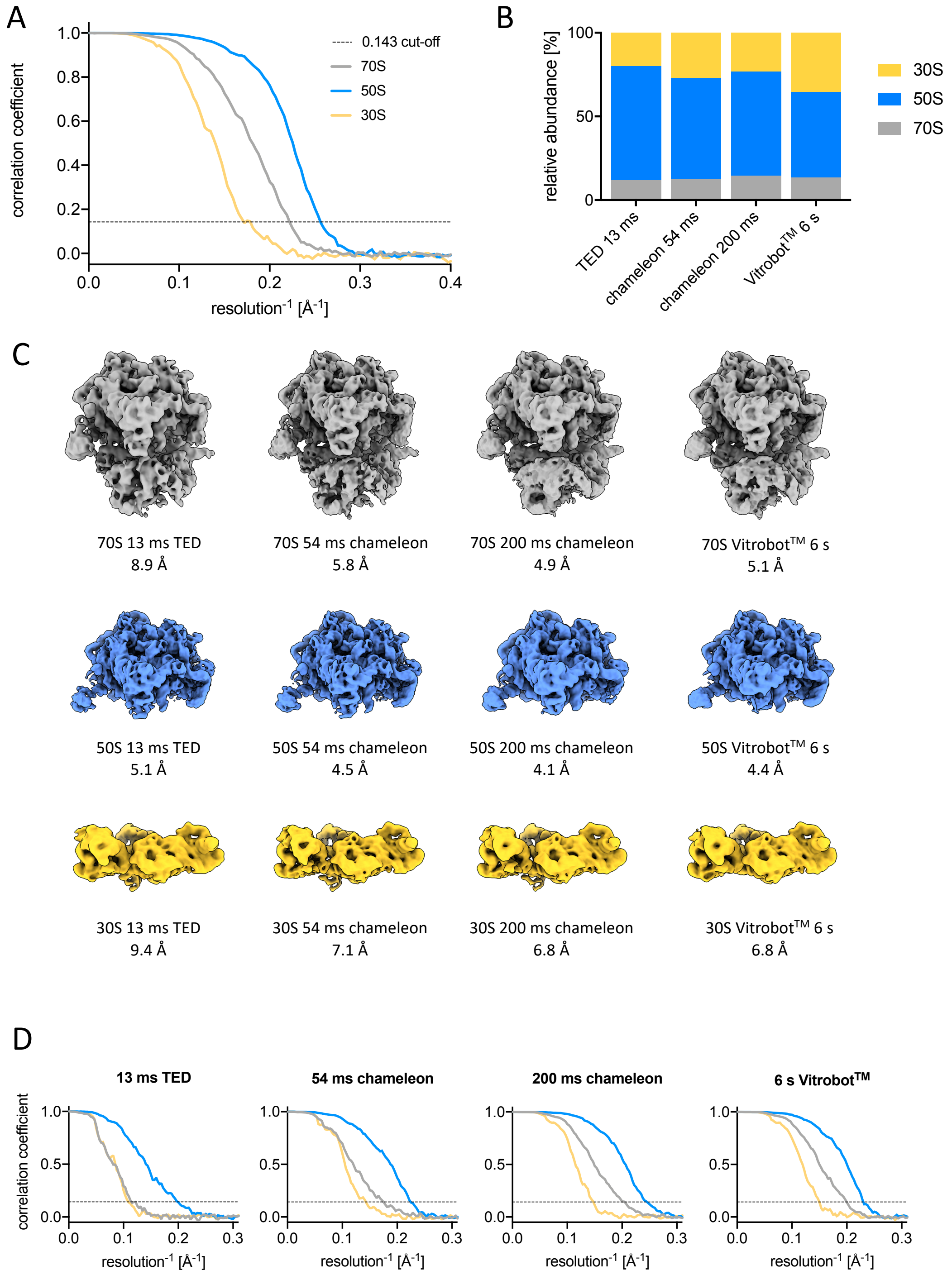


**CryoEM reconstructions of ribosome data at different time points and vitrification devices.** (A) FSC curves for masked, consensus reconstructions of 70S, 50S and 30S ribosome. (B) Relative particle numbers for 30S, 50S and 70S by individual dataset. (C) Individual reconstructed maps for 70S, 50S and 30S for all 4 subsets and corresponding FSC curves (D).

**Table S1.**

| | Time and vitrification device | Repeat | Concentration [µM] | Applied concentration [µM] | Ice thickness [nm] | # particles | relative particle # (≤ 10nm) | relative particle # (≤ 20nm) |
|---|---|---|---|---|---|---|---|---|
| Apoferritin | 11 ms TED | 1 | 1.0 | 20.0 | 168 | 100 | 21* | 61 |
| | | 2 | 1.9 | | 96 | 111 | 77 | 86 |
| | | 3 | 1.4 | | 78 | 64 | 73 | 89 |
| | | 4 | 0.9 | | 97 | 53 | 74 | 91 |
| | | 5 | 2.2 | | 139 | 186 | 75 | 89 |
| | 50 ms TED | 1 | 3.4 | 20.0 | 152 | 253 | 72 | 77 |
| | | 2 | 7.7 | | 135 | 509 | 66 | 74 |
| | | 3 | 2.1 | | 68 | 54 | 48 | 72 |
| | | 4 | 1.7 | | 76 | 79 | 63 | 82 |
| | | 5 | 1.4 | | 64 | 54 | 59 | 76 |
| | | 6 | 119.2 | | 47 | 2160 | 90 | 98 |
| | | 7 | 41.8 | | 90 | 1835 | 91 | 97 |
| | | 8 | 1.3 | | 88 | 54 | 46 | 52 |
| | 6 s Vitrobot™ | 1 | 53.9 | 20.0 | 54 | 1751 | 90 | 97** |
| | | 2 | 69.1 | | 43 | 1622 | 82 | 96** |
| | | 3 | 83.8 | | 34 | 1582 | 87 | 99** |
| HSPD1 | 6 ms TED | 1 | 1.5 | 11.0 | 95 | 70 | 77 | 87 |
| | | 2 | 1.2 | | 73 | 69 | 46* | 81 |
| | | 3 | 0.9 | | 82 | 37 | 97 | 97 |
| | | 4 | 2.3 | | 62 | 85 | 93 | 96 |
| | 50 ms TED | 1 | 18.8 | 11.0 | 60 | 680 | 99 | 100 |
| | | 2 | 4.9 | | 87 | 204 | 85 | 87 |
| | | 3 | 7.4 | | 73 | 325 | 100 | 100 |
| | 6 s Vitrobot™ | 1 | 18.9 | 0.6 | 39 | 193 | 100 | 100** |
| | | 2 | 10.0 | | 50 | 270 | 99 | 100** |
| | | 3 | 9.3 | | 64 | 156 | 98 | 100** |
| Ribosome | 13 ms TED | 1 | 1.3 | 2.5 | 113 | 57 | 19* | 100 |
| | | 2 | 0.3 | | 130 | 14 | 100 | 100 |
| | | 3 | 3.2 | | 151 | 84 | 94 | 98 |
| | | 4 | 3.9 | | 119 | 178 | 92 | 99 |
| | | 5 | 5.9 | | 67 | 152 | 99 | 99 |
| | 200 ms chameleon | 1 | 11.9 | 2.5 | 136 | 864 | 95 | 96 |
| | | 2 | 12.5 | | 137 | 934 | 94 | 96 |
| | | 3 | 15.8 | | 135 | 1235 | 92 | 96 |
| | 6 s Vitrobot™ | 1 | 17.0 | 0.8 | 92 | 601 | 79 | 84 |
| | | 2 | 23.5 | | 93 | 843 | 84 | 87 |
| | | 3 | 20.2 | | 97 | 787 | 78 | 83 |

**Summary table of tomograms analysed to produce partitioning and particle concentration data.** "Concentration" is the estimated concentration from the tomogram, "Applied concentration" is the concentration of the sample applied. "Relative particle #" is the percentage of particles within ≤ 10 or 20 nm of the AWI.

* Values were excluded from analysis because of poor fitting of AWIs.

** Values were excluded from analysis because of low ice thickness.

Table S2. Microscope parameters for collection of cryo-ET data

| Microscope | Titan Krios II |
|---|---|
| Magnification | 42,000 |
| Voltage (kV) | 300 |
| Electron dose per image ($e^-/Å^2$) | 1.8 - 1.9 |
| Fractions per image | 3 |
| maximum/minimum tilt (increment) | -60°, +60° (2°) |
| Tilt scheme | bidirectional |
| Defocus range (µm) | -5 to -10 |
| Pixel size (Å) | 3.4 |

Table S3. Data collection parameters for SPA datasets of HSPD1.

| | **HSPD1** | | | |
|---|---|---|---|---|
| | **6 ms TED** | **50 ms TED** | **54 ms chameleon** | **6 s Vitrobot™** |
| Microscope | Titan Krios I | | | |
| Magnification | 75,000 | | | |
| Voltage (kV) | 300 | | | |
| Total electron dose ($e^-/Å^2$) | 81 | 81 | 74 | 75 |
| Exposure time | 1.5 | 1.5 | 1.5 | 1.5 |
| Number of frames | 59 | 59 | 59 | 59 |
| Defocus range (µm) | -2 to -4 | -2 to -4.5 | -1.5 to -3.5 | -1.3 to -3.3 |
| Pixel size (Å) | 1.065 | | | |

**Table S4**. Data collection parameters for SPA datasets of ribosome.

| | **ribosome** | | | |
|---|---|---|---|---|
| | **13 ms TED** | **54 ms chameleon** | **200 ms chameleon** | **6 s Vitrobot™** |
| Microscope | Titan Krios I | | | |
| Magnification | 75,000 | | | |
| Voltage (kV) | 300 | | | |
| Total electron dose ($e^-/Å^2$) | 77 | 74 | 74 | 78 |
| Exposure time | 1.5 | 1.6 | 1.5 | 1.5 |
| Number of frames | 59 | 59 | 59 | 59 |
| Defocus range (µm) | -1.3 to -3.3 | -1.3 to -3.3 | -1.3 to -3.3 | -1.3 to -3.3 |
| Pixel size (Å) | 1.065 | | | |
